# ANTIBACTERIAL ACTIVITY OF LEAVES AND STEM OF *Bryohyllum Pinnatum* (MIRACLE LEAF) IN SOKOTO STATE, NIGERIA

**DOI:** 10.1101/2025.02.09.637277

**Authors:** Maryam Dantani, Amina Musa Rabe, Abdulmalik Dantani

## Abstract

This study aimed to evaluate the antibacterial activity of *Bryophyllum pinnatum* leaves and stems against selected bacterial isolates from Sokoto State, Nigeria. Aqueous and ethanol solvents were used to extract bioactive compounds, and the extracts were tested at concentrations of 200, 140, and 60 mg/ml against *Staphylococcus aureus, Pseudomonas aeruginosa*, and *Micrococcus spp*. The results revealed no inhibitory effects from the *Bryophyllum pinnatum* extracts on the tested bacterial strains, irrespective of solvent type or concentration. In contrast, the standard antibiotic Tetracycline exhibited significant antibacterial activity, with inhibition zones of 20 mm, 16 mm, and 9 mm for *Staphylococcus aureus, Micrococcus spp*., and *Pseudomonas aeruginosa*, respectively.

These findings suggest that *Bryophyllum pinnatum* extracts, under the tested conditions, lack antibacterial activity against the evaluated bacterial isolates. However, this does not exclude the plant’s potential for other therapeutic applications. Future studies should investigate alternative extraction methods, diverse solvent systems, and a broader range of bacterial strains to comprehensively evaluate the bioactivity of *Bryophyllum pinnatum*.

## INTRODUCTION

In recent years, the rise of antibiotic-resistant strains among clinically significant pathogens has become a pressing concern, leading to the emergence of multi-resistant bacterial strains [1,2,3]. The scarcity and high cost of new-generation antibiotics, coupled with their limited effectiveness, have contributed to increased rates of morbidity and mortality [4]. Consequently, there is a growing imperative to explore alternative antimicrobial substances from various sources. This has prompted investigations into the antimicrobial properties of materials derived from plants, aiming to identify potentially potent active compounds that could serve as the basis for new antimicrobial drugs [5,6].

*Bryophyllum pinnatum* (*Kalanchoe pinnata*; Lamarch Crassulaceae) is an erect, succulent, perennial shrub that grows about 1.5m tall and reproduces through seeds and also vegetatively from leaf bulbils [7]. It has a tall hollow stems, freshly dark green leaves that are distinctively scalloped and trimmed in red and dark bell-like pendulous flowers [8]. *Bryophyllum pinnatum* can be propagated through stems or leaf cutting. It is an introduced ornamental plant that is now growing as a weed around plantation crop [7,9]. *Bryophyllum pinnatum* is used in ethnomedicine for the treatment of earache, burns, abscesses, ulcers, insect bites, whitlow, diarrhea and cithiasis [9]. In Southeastern Nigeria, this herb is used to facilitate the dropping of the placenta of new born baby [9]. The lightly roasted leaves are used externally for skin fungus and inflammations. The leaf infusion is an internal remedy for fever [8].

*Bryophyllum pinnatum* holds a prominent place in ethnomedicine for its diverse applications, including the treatment of earaches, burns, ulcers, insect bites, diarrhea, and postpartum issues like placental expulsion [9]. It is valued for its sedative, wound-healing, diuretic, anti-inflammatory, and cough-suppressant properties, making it a versatile remedy for a range of health issues [8,10]. In ethnomedicine, *Bryophyllum pinnatum* is used to induce vomiting of blood, cut, umbilical cord in new born baby, expel worms, cure acute and chronic bronchitis, pneumonia and other forms of respiratory tract disease. The plant is regarded as a sedative wound-healer, diuretic, anti-inflammatory and cough suppressant [8]. It is used for the treatment of all sorts of respiratory ailments such as asthma, cough and bronchitis [8]. It is employed for the treatment of kidney stones, gastric ulcers and edema of the leg [10]. The juice from the leaf is used to treat boils and skin ulcers. *Bryophyllum pinnatum* is widely used in herbal medicine to cure diseases and heal injuries. Despite its widespread use in traditional medicine, limited research has rigorously evaluated its antibacterial properties.

Nigeria is richly endowed with indigenous plants used in herbal medicine to treat diseases and injuries [7]. Many of these plants exhibit diverse biological and pharmacological activities, including anti-inflammatory, diuretic, antispasmodic, antihypertensive, antidiabetic, and antimicrobial functions. Medicinal plants represent a rich reservoir of novel drug compounds that contribute to traditional medicine systems, modern pharmaceuticals, nutraceuticals, and folk remedies [11]. The World Health Organization (WHO) has highlighted that over 80% of the global population relies on plants to fulfill their primary healthcare needs [11]. With increasing demand driving cultivation efforts, medicinal plants like *Bryophyllum pinnatum* offer the potential to bridge the gap between traditional knowledge and modern scientific applications [12].

The aim of this study is to evaluate the antibacterial properties of *Bryophyllum pinnatum* leaves and stems, focusing on its potential to address the growing challenge of antibiotic resistance. Specifically, this research investigates the antibacterial activity of aqueous and ethanol extracts of *Bryophyllum pinnatum* against clinically significant bacterial strains, including *Staphylococcus aureus, Pseudomonas aeruginosa*, and *Micrococcus* spp.

By building on the ethnomedicinal significance of *Bryophyllum pinnatum*, this study seeks to contribute to the scientific understanding of its potential as a *source* of new antimicrobial agents. Additionally, it highlights the importance of validating traditional remedies through rigorous scientific methods, aiming to address the global challenge of antibiotic resistance while supporting the development of plant-based therapeutic alternatives.

While *Bryophyllum pinnatum* has a long history of use in traditional medicine, its potential antibacterial properties remain underexplored, particularly against pathogens contributing to healthcare-associated infections such as *Staphylococcus aureus* and *Pseudomonas aeruginosa*. This study aims to bridge this knowledge gap, providing insights that could lead to the development of novel, affordable plant-based antibacterial therapies.

## MATERIALS AND METHODS

### Study Area

The study was conducted at the Mycology and Microbiology Laboratories of Usmanu Danfodiyo University, Sokoto. Sokoto is located at the extreme northwest of Nigeria, near the confluence of the River Sokoto and the River Rima (Latitude 13°N, Longitude 5°E, and 350m above sea level). This region is characterized by a semi-arid climate, unique soil composition, and specific vegetation types, which may influence the chemical properties and bioactivity of plants. Research conducted in such a geographical area is vital for understanding how local environmental factors, such as climate and soil nutrients, impact the medicinal potential of *Bryophyllum pinnatum*. This ensures the findings are relevant to both the local population and broader scientific understanding.

### Collection and Identification of Plant Materials

Fresh and mature leaves and stems of *Bryophyllum pinnatum* (commonly known as the Miracle Leaf) were collected in May 2023 from the Biological Garden of Usmanu Danfodiyo University, Sokoto, Nigeria. The plant was authenticated and identified by a taxonomist at the Herbarium of the Plant Science Department, Usmanu Danfodiyo University, Sokoto. Authentication of the plant material is crucial to ensure the validity and reproducibility of the study, as misidentification could compromise the reliability of the findings.

### Preparation of Plant Material

The collected leaves and stems were separated and thoroughly washed with tap water to remove dust and debris. The leaves and stems were then air-dried at room temperature, with the stems requiring 3 weeks and the leaves 4 weeks to dry due to their differences in thickness and architecture. To preserve the active compounds, the leaves were not exposed to direct sunlight, following the method described by Girish and Satish (2000) [13].

Once dried, the leaves and stems were pulverized into a coarse powder using a mortar and pestle. This manual grinding method was chosen over mechanical grinders to minimize heat generation, which could degrade sensitive bioactive compounds. The powdered material was then sieved using a 2mm mesh (British standard). The mesh size was selected to ensure uniform particle size, which facilitates efficient solvent penetration during the extraction process. The fine powdered leaves and stems were stored in sterile, airtight containers until needed for extraction and further analysis.

### Collection of Experimental Microorganisms

The bacterial strains used in this study—Staphylococcus aureus, Pseudomonas aeruginosa, and Micrococcus spp.—were laboratory strains obtained from the Microbiology Laboratories of Usmanu Danfodiyo University, Sokoto. These isolates were sub-cultured onto nutrient agar slants and incubated at 37°C for 24 hours to ensure pure and viable cultures for the experiments.

The selected bacterial strains are commonly used in antimicrobial studies due to their well-documented characteristics and relevance in laboratory testing. Staphylococcus aureus is frequently studied for its role in infections and antibiotic resistance. Pseudomonas aeruginosa is an opportunistic pathogen known for its multidrug-resistant properties, and Micrococcus spp. serves as a model organism for testing antimicrobial activity due to its ubiquitous presence and ease of handling. The use of these laboratory strains ensures reproducibility and standardization in evaluating the antibacterial potential of *Bryophyllum pinnatum* extracts.

### Preparation of Aqueous and Ethanolic Extracts

The extraction methods described by Harborne (1973) and Aliyu et al. (2009) were adopted. A total of 100 g of the powdered plant material (leaves and stem) was soaked in 1 liter of distilled water and ethanol, respectively. The choice of these solvents was based on their polarity differences, allowing for the extraction of a broad range of bioactive compounds. Water, being a polar solvent, is effective for extracting hydrophilic compounds, while ethanol, a moderately polar solvent, facilitates the extraction of both hydrophilic and lipophilic compounds [14, 15].

The mixtures were left overnight for 24 hours to ensure thorough extraction, then decanted and filtered to remove solid residues. The filtrates were evaporated in a hot dry oven set at 40°C to prevent the degradation of heat-sensitive compounds. The resulting residues were carefully collected and stored in sterile containers at room temperature until required for sensitivity testing.

### Media preparation

Nutrient Agar media was prepared following the manufacturer’s instructions. A total of 28 g of the powder was measured and dissolved in 1000 ml of distilled water in a 2000 ml conical flask. The mixture was stirred thoroughly to ensure complete dissolution and then covered with cotton wool and aluminum foil. The media was sterilized by autoclaving at 121°C for 15 minutes, a crucial step to eliminate potential contaminants.

After sterilization, small amounts of the media were poured into Petri dishes. The media was allowed to cool and solidify, forming the base layer. An additional layer of media was poured on top and allowed to cool at room temperature, creating a two-layered structure. This layering method was used to enhance even distribution of the extracts and facilitate better diffusion during the antibacterial activity test.

### Screening for antibacteral activity

The antibacterial activity of *Bryophyllum pinnatum* extracts was evaluated using the agar well diffusion method described by Atata et al. (2003). A bacterial suspension was prepared in sterile water and standardized to match the turbidity of a 0.5 McFarland standard, ensuring a uniform bacterial concentration [16]. The tested bacterial isolates included Staphylococcus aureus, Pseudomonas aeruginosa, and Micrococcus spp.

Each nutrient agar plate was uniformly inoculated with the standardized bacterial suspension using a sterile cotton swab. Wells of 6 mm diameter were created in the inoculated agar medium using a sterile cork borer, which provided consistent well sizes for reproducibility and optimal diffusion of the extracts. A volume of 0.1 ml of each *Bryophyllum pinnatum* extract (aqueous and ethanolic) was introduced into the wells. Tetracycline, a widely used antibiotic, was used as a positive control.

The plates were left to stand at room temperature for 30 minutes to allow initial diffusion of the extracts into the medium. Subsequently, the plates were incubated at 37°C for 24 hours to facilitate bacterial growth and the evaluation of antibacterial activity. Zones of inhibition, indicating areas where bacterial growth was suppressed, were measured in millimeters using a ruler. All experiments were performed in triplicates to ensure statistical reliability and consistency of the results.

## Results

Table 1 summarizes the antibacterial activity of the ethanolic and aqueous extracts of *Bryophyllum pinnatum* leaves and stems against the tested microorganisms: *Staphylococcus aureus (S. aureus), Micrococcus spp (M. spp)*, and *Pseudomonas aeruginosa (P. aeruginosa)*.

**Table 1.** Antibacterial activity of Stem and leaves extract of *Bryophyllum pinnatum* on test microorganisms.

| Extract | Bacteria |  |  |  |
| --- | --- | --- | --- | --- |
|  | Concentration (mg/ml) | *S. aureus* | *M. spp* | *P. aeruginosa* |
| Ethanolic extract | 60 | 0.00 | 0.00 | 0.00 |
|  | 140 | 0.00 | 0.00 | 0.00 |
|  | 200 | 0.00 | 0.00 | 0.00 |
| Aqueous extract | 60 | 0.00 | 0.00 | 0.00 |
|  | 120 | 0.00 | 0.00 | 0.00 |
|  | 200 | 0.00 | 0.00 | 0.00 |
| Control (Tetracycline) | 10µg | 20.0 | 16.0 | 9.00 |

0.00 = No observable inhibitory effect (zone of inhibition)

At concentrations of 60 mg/ml, 140 mg/ml, and 200 mg/ml, neither the ethanolic nor the aqueous extracts exhibited any inhibitory effect on the growth of the tested bacteria. The absence of inhibitory activity is denoted by “0.00,” indicating no measurable zones of inhibition.

In contrast, the control antibiotic, tetracycline, at a concentration of 10 μg, demonstrated significant inhibitory effects, with measurable zones of inhibition against all tested bacteria. The zones were recorded as 20.0 mm for *S. aureus*, 16.0 mm for *M. spp*, and 9.0 mm for *P. aeruginosa*. This highlights the control’s effectiveness compared to the plant extracts under study.

## Discussion

The use of plants for medicinal purposes is an age-old tradition in Africa, Asia, and Latin America [17,18]. The effectiveness of these medicinal plants is repeatedly validated in the laboratory, with roughly 80% of plants selected for analysis based on ethnomedicinal information demonstrating significant pharmacological activity [19]. However, in this study, the ethanolic and aqueous extracts of *Bryophyllum pinnatum* (leaves and stem) did not inhibit the growth of *Staphylococcus aureus, Pseudomonas aeruginosa*, and *Micrococcus spp*, whereas the control drug, tetracycline, effectively inhibited their growth. This result may be attributed to several factors, including the bacteria’s strong immune systems, extraction solvents, or experimental methodology.

The inability of *Bryophyllum pinnatum* extracts to inhibit bacterial growth aligns with findings from previous studies. For instance, Mukhtar and Tukur (2001) noted that *Pseudomonas aeruginosa* is particularly resistant, with reports suggesting that this bacterium had developed resistance to many antibiotics even before their discovery [20]. Similarly, Faleye and Ogundaini reported that extracts of *Aspilia africana* obtained using six different solvents showed no activity against *P. Aeruginosa* [21]. These findings suggest that the choice of solvent plays a critical role in extracting active antimicrobial agents.

Despite the lack of observed antimicrobial activity in this study, the ethnomedicinal uses of *Bryophyllum pinnatum* have been partly rationalized by previous work. For example, Oladejo et al. (2021) reported that ethanol extracts of *Bryophyllum pinnatum* inhibited the growth of *P. aeruginosa* at a lower concentration of 15.625 mg/ml, achieving a zone of inhibition of 15.00 ± 0.02 mm [22]. This highlights the potential concentration-dependent nature of the plant’s antimicrobial properties, where lower concentrations may improve solubility and efficacy of active compounds.

The choice of solvents (ethanol and aqueous) in this study may have limited the extraction of active antimicrobial agents. Akinpelu (2000) and Ofokansi (2005) reported that methanol extracts of *Bryophyllum pinnatum* exhibited strong activity against some Gram-positive organisms, likely due to methanol’s superior ability to extract antimicrobial compounds such as phenolic compounds, saponins, bryophyllins, and other secondary metabolites This suggests that the lack of antimicrobial activity in this study may be due to the solvents’ inability to dissolve certain active principles [23, 24]. Furthermore, Ellof (1998) noted that improper drying techniques might result in the loss of active compounds, further contributing to the lack of observed activity [25].

While this study’s results reveal no antimicrobial activity of *Bryophyllum pinnatum* extracts against the tested bacterial strains, it underscores the complexity of plant extracts and their interactions with microorganisms. Ethnomedicinal uses of the plant may extend beyond antimicrobial activity, encompassing anti-inflammatory, wound-healing, or other therapeutic effects. This highlights the need to integrate traditional knowledge with systematic research to explore the plant’s full pharmacological potential.

To address the limitations observed in this study, future research should Explore the use of methanol or other solvents with better extraction capabilities, optimize extraction protocols, including solvent concentrations and drying techniques, to preserve active compounds. Test a wider range of concentrations, particularly lower levels, to examine concentration-dependent efficacy. Investigate the plant’s activity against a broader spectrum of microorganisms, including fungi and other bacteria. Conduct phytochemical analyses to identify specific active compounds responsible for the plant’s pharmacological properties.

This study emphasizes the importance of extraction methods, solvent choice, and experimental design in determining the antimicrobial potential of plant extracts. While the results indicate no activity of *Bryophyllum pinnatum* against the tested bacteria at the tested concentrations, they provide a foundation for refining methodologies and exploring alternative approaches to uncover the plant’s potential benefits.

## Conclusion

This study demonstrated that Bryophyllum pinnatum extracts from stems and leaves, prepared using ethanol and aqueous solvents, exhibited no antibacterial activity against *Staphylococcus aureus, Pseudomonas aeruginosa*, and *Micrococcus spp*. at concentrations of 60, 140, and 200 mg/ml. Further studies are recommended to investigate the antibacterial potential of *B. pinnatum* using alternative solvents and optimized extraction methods to isolate active antimicrobial compounds and validate these findings.

